## Supplementary material for "A single-cell atlas of spatial and temporal gene expression in the mouse cranial neural plate": Figure Supplements

### Figure 1 - Figure Supplement 1

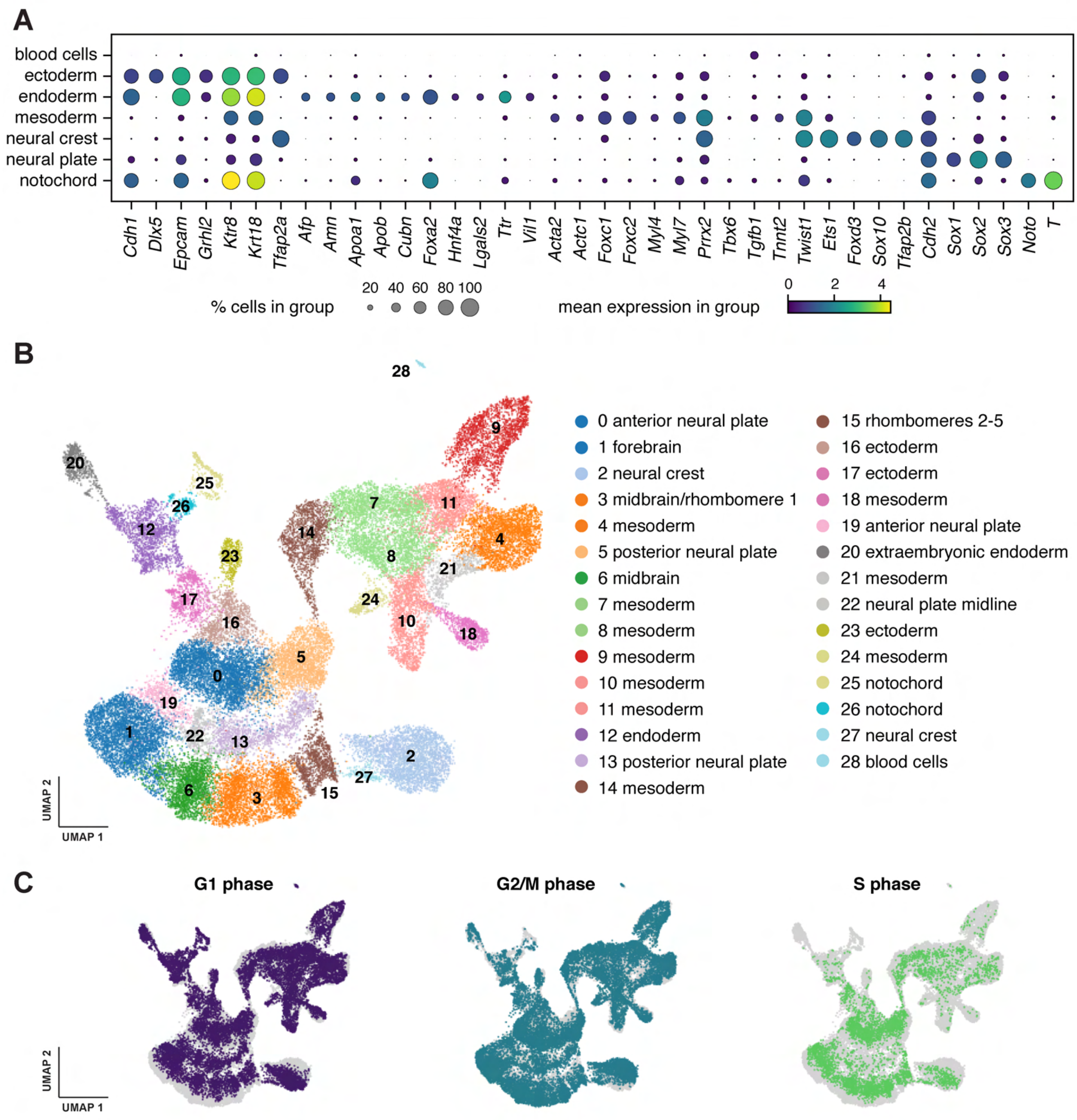

#### Figure 1 - Figure Supplement 2

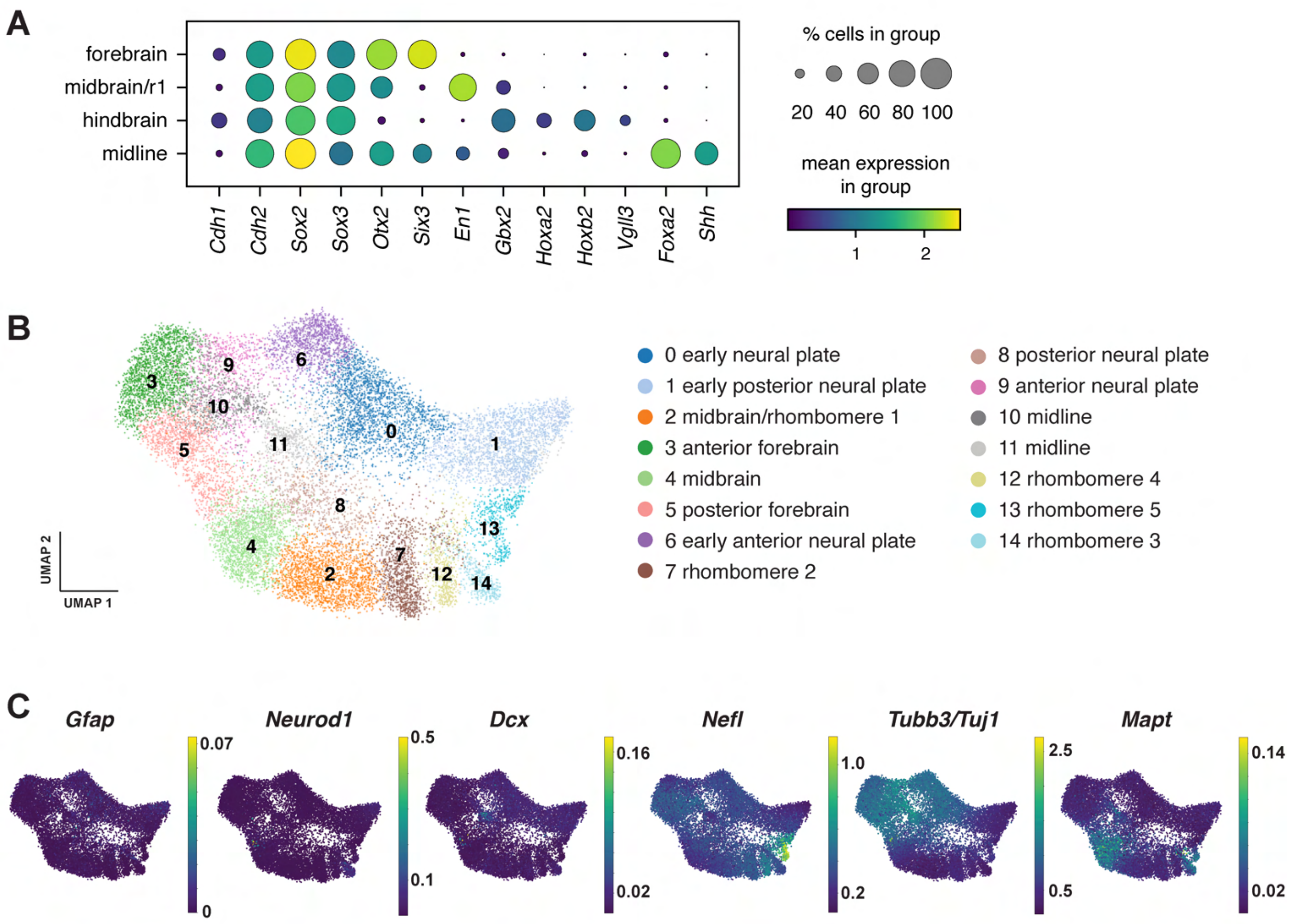

### Figure 2 - Figure Supplement 1

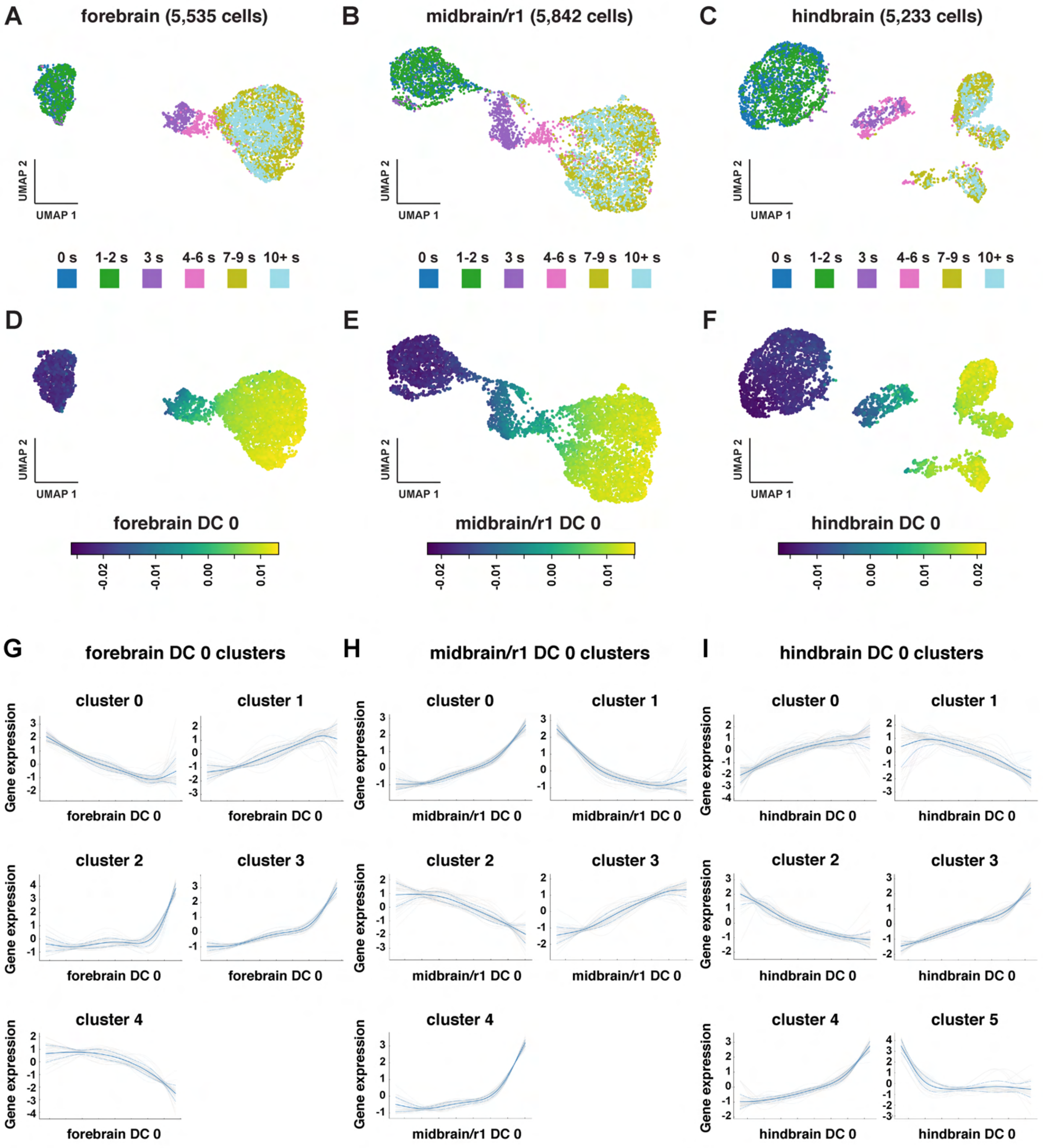

**Figure 2 - Figure Supplement 2**

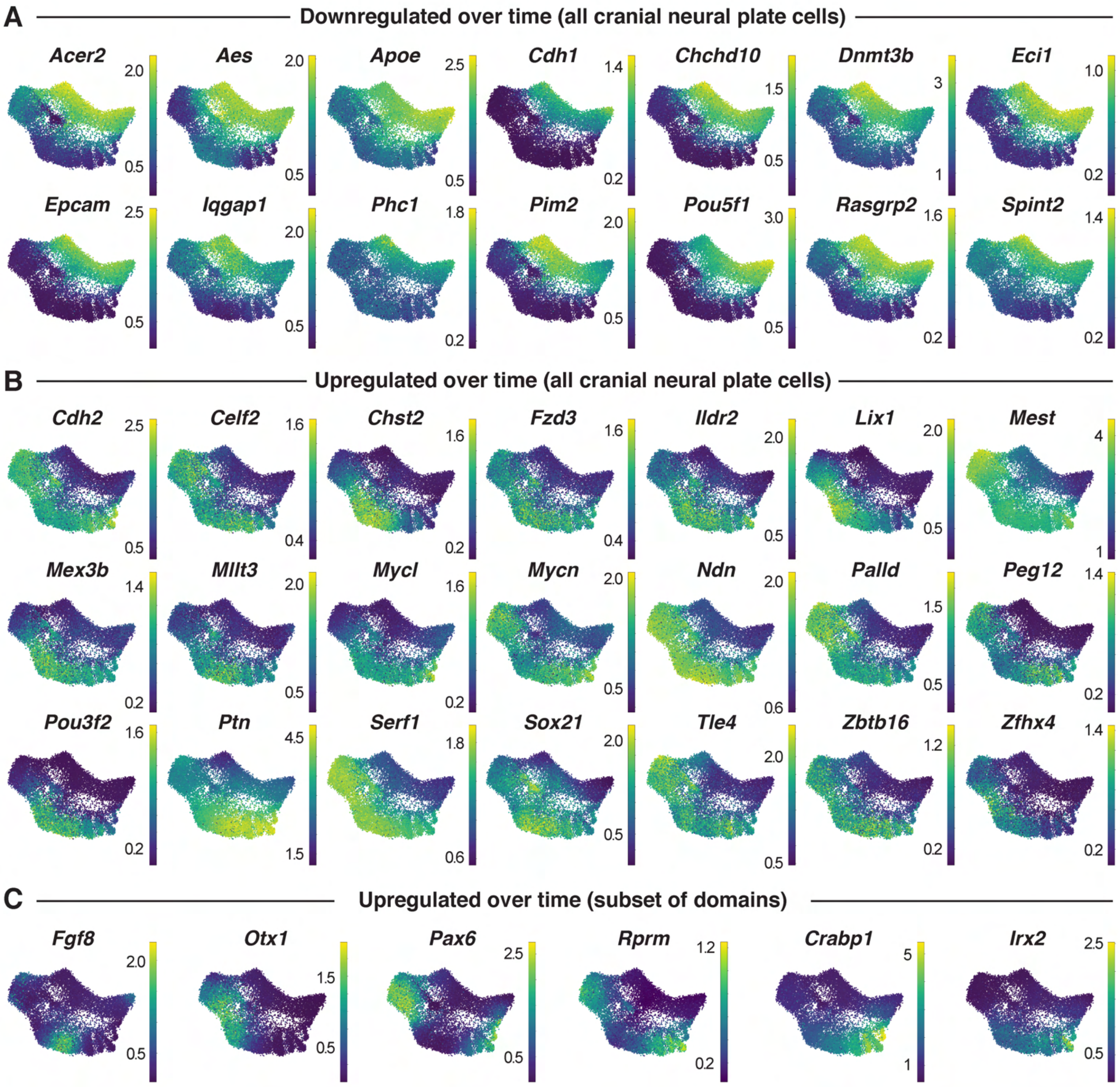

### Figure 2 - Figure Supplement 3

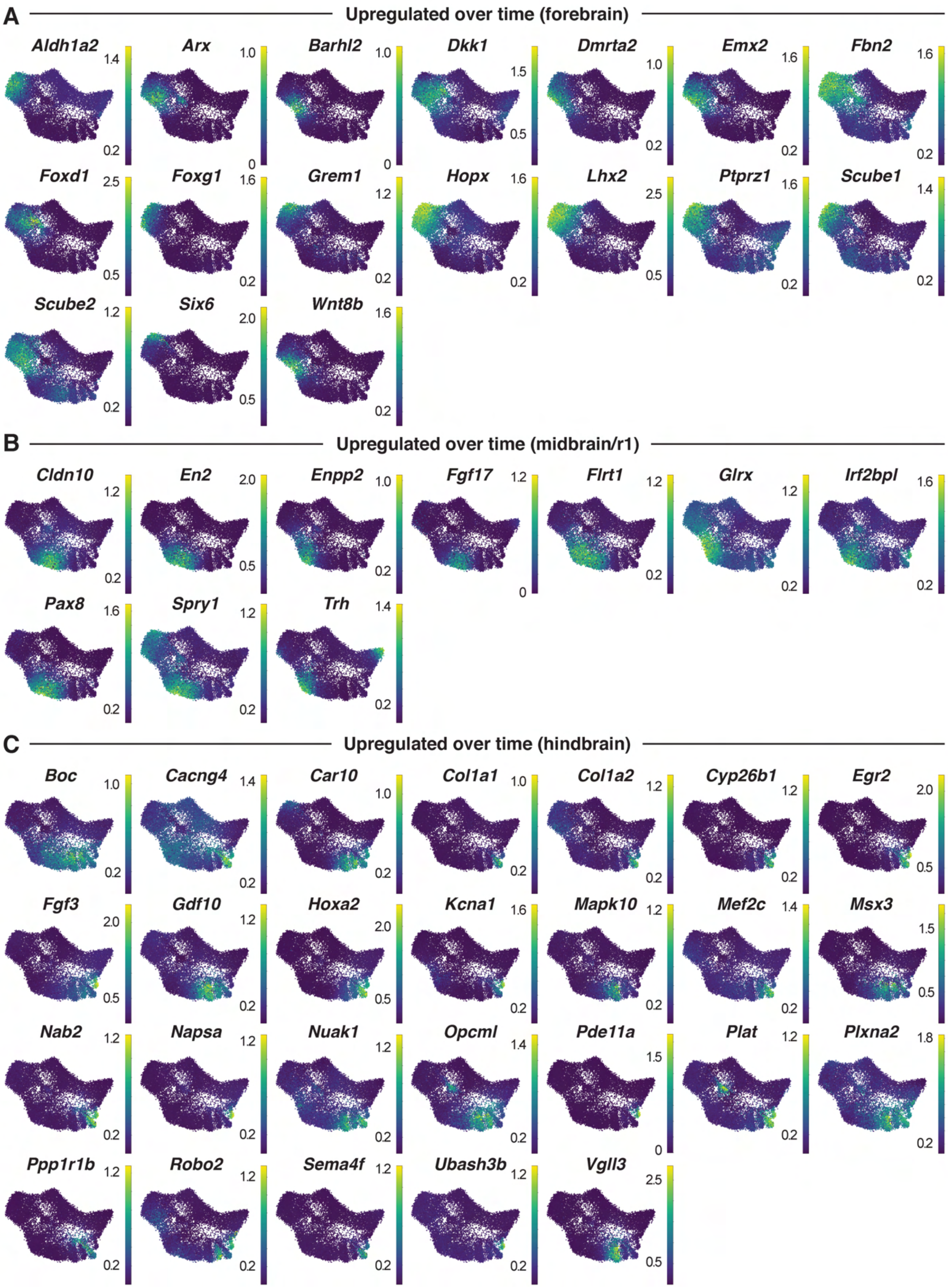

### Figure 3 - Figure Supplement 1

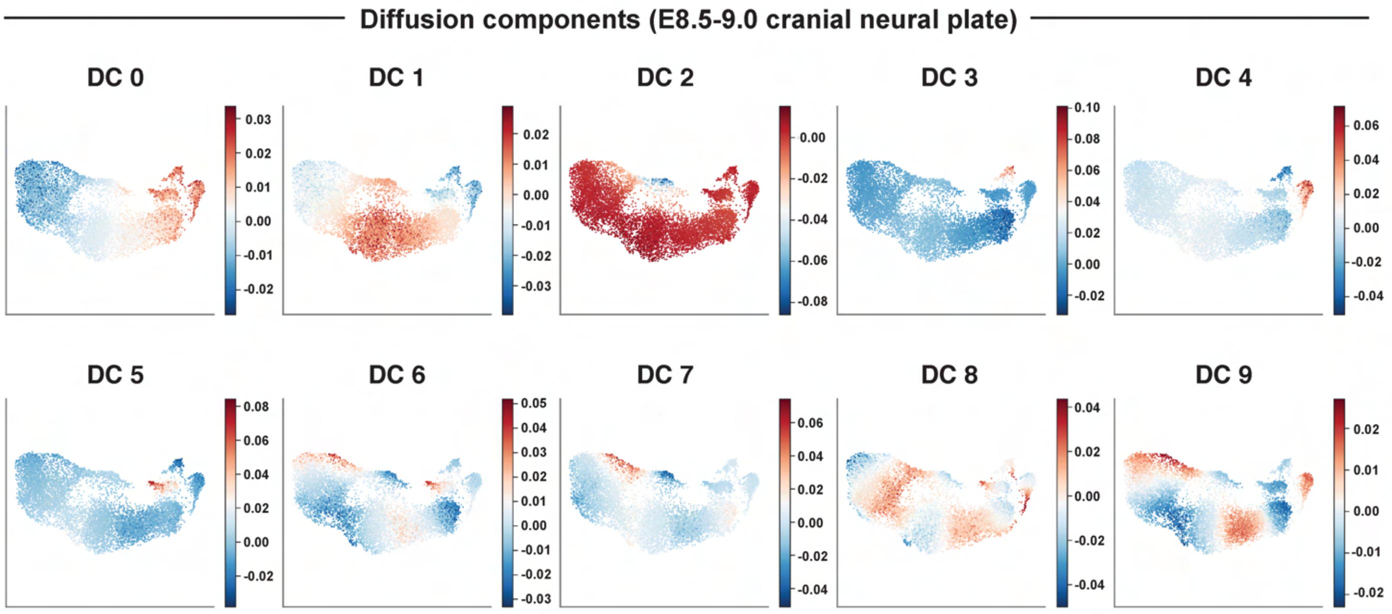

### Figure 3 - Figure Supplement 2

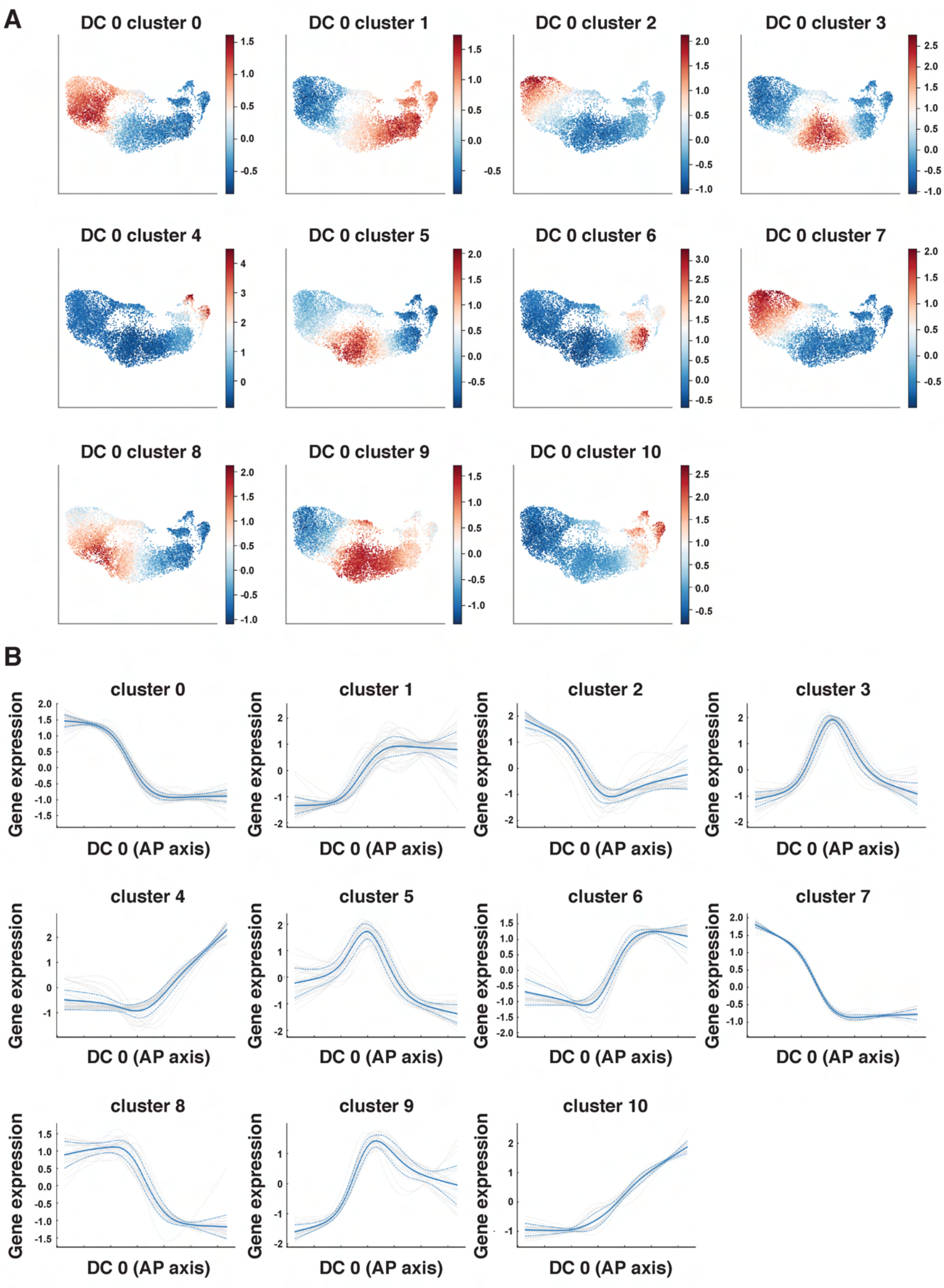

### Figure 4 - Figure Supplement 1

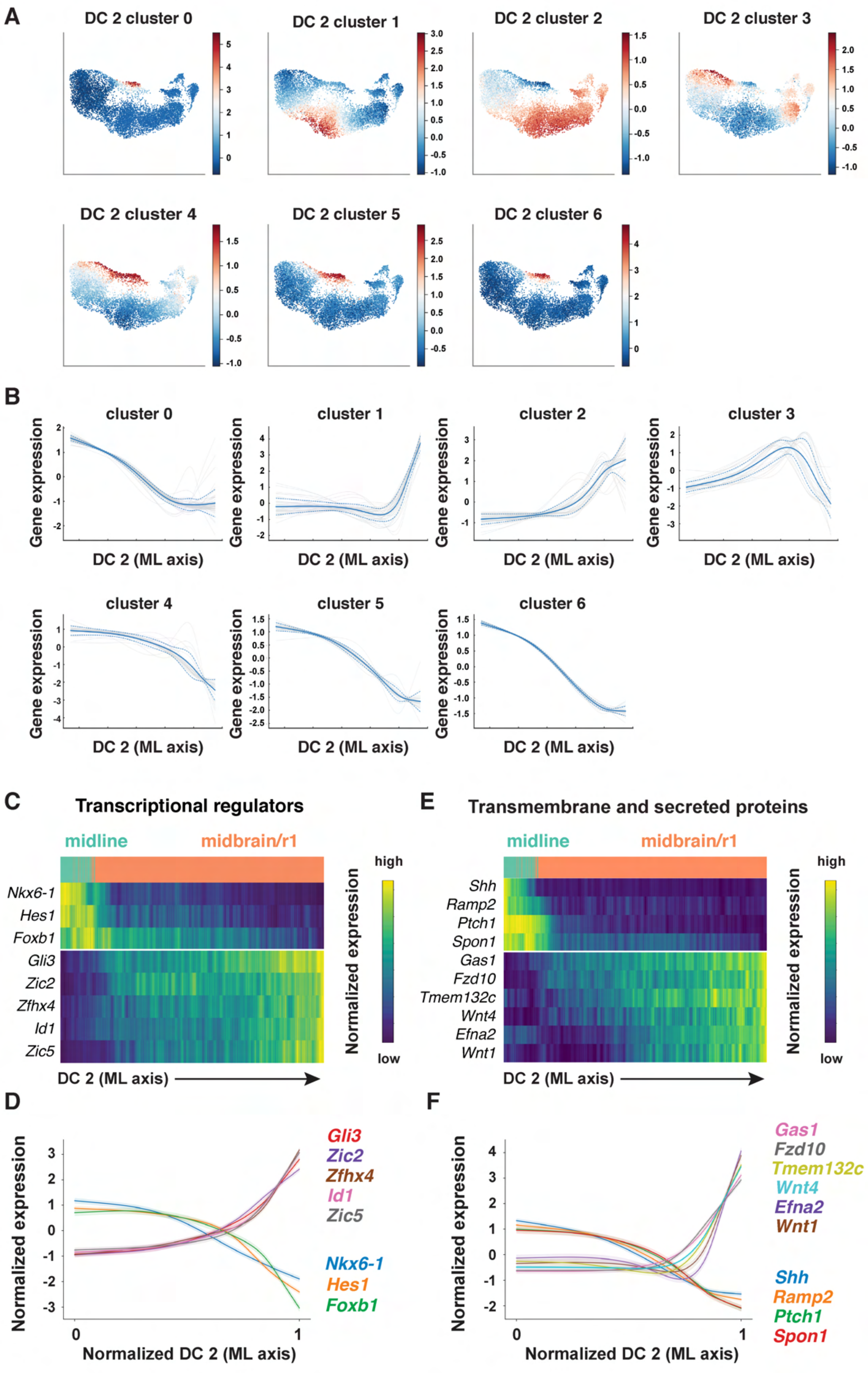

#### Figure 5 - Figure Supplement 1

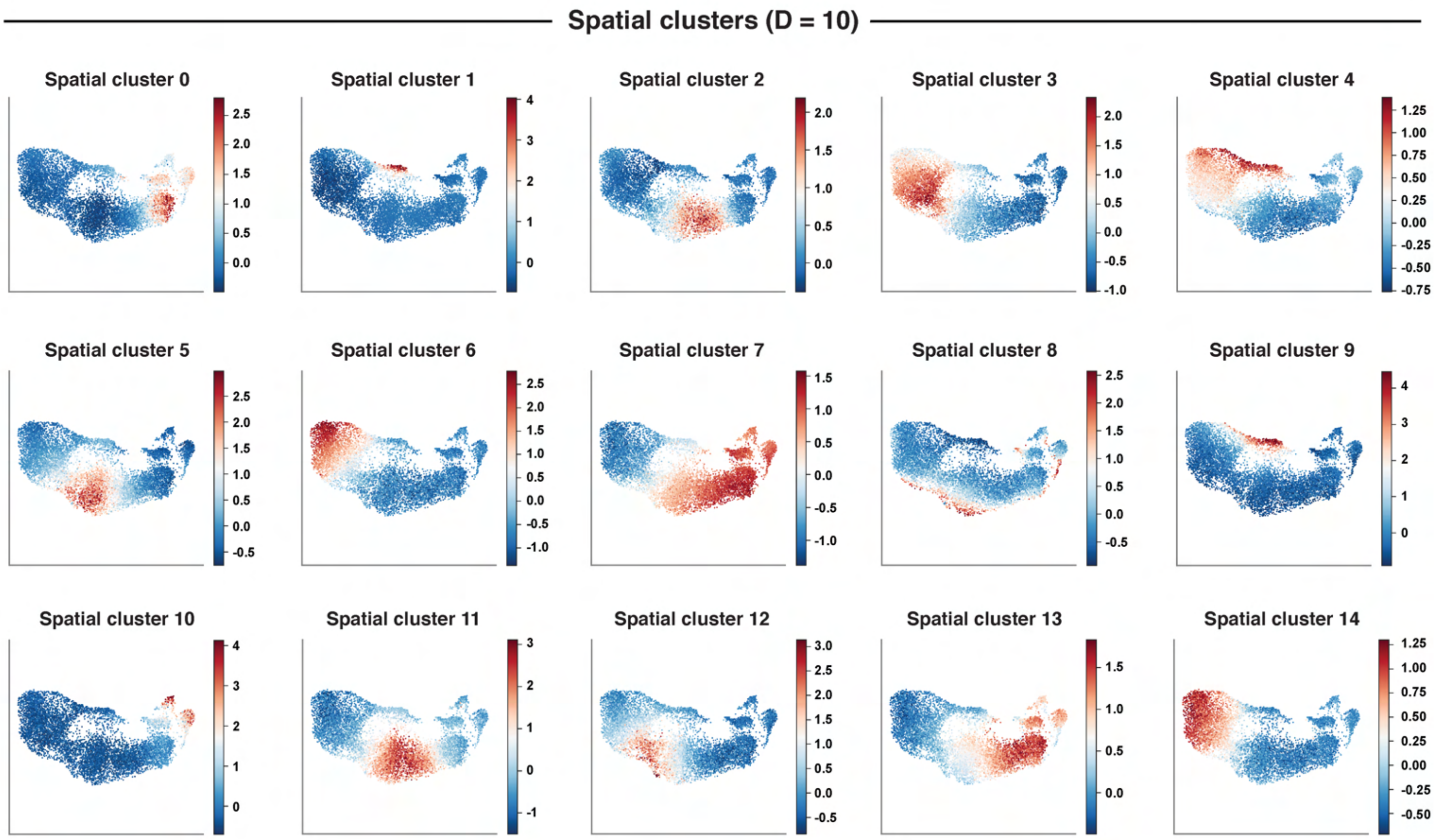

### Figure 5 - Figure Supplement 2

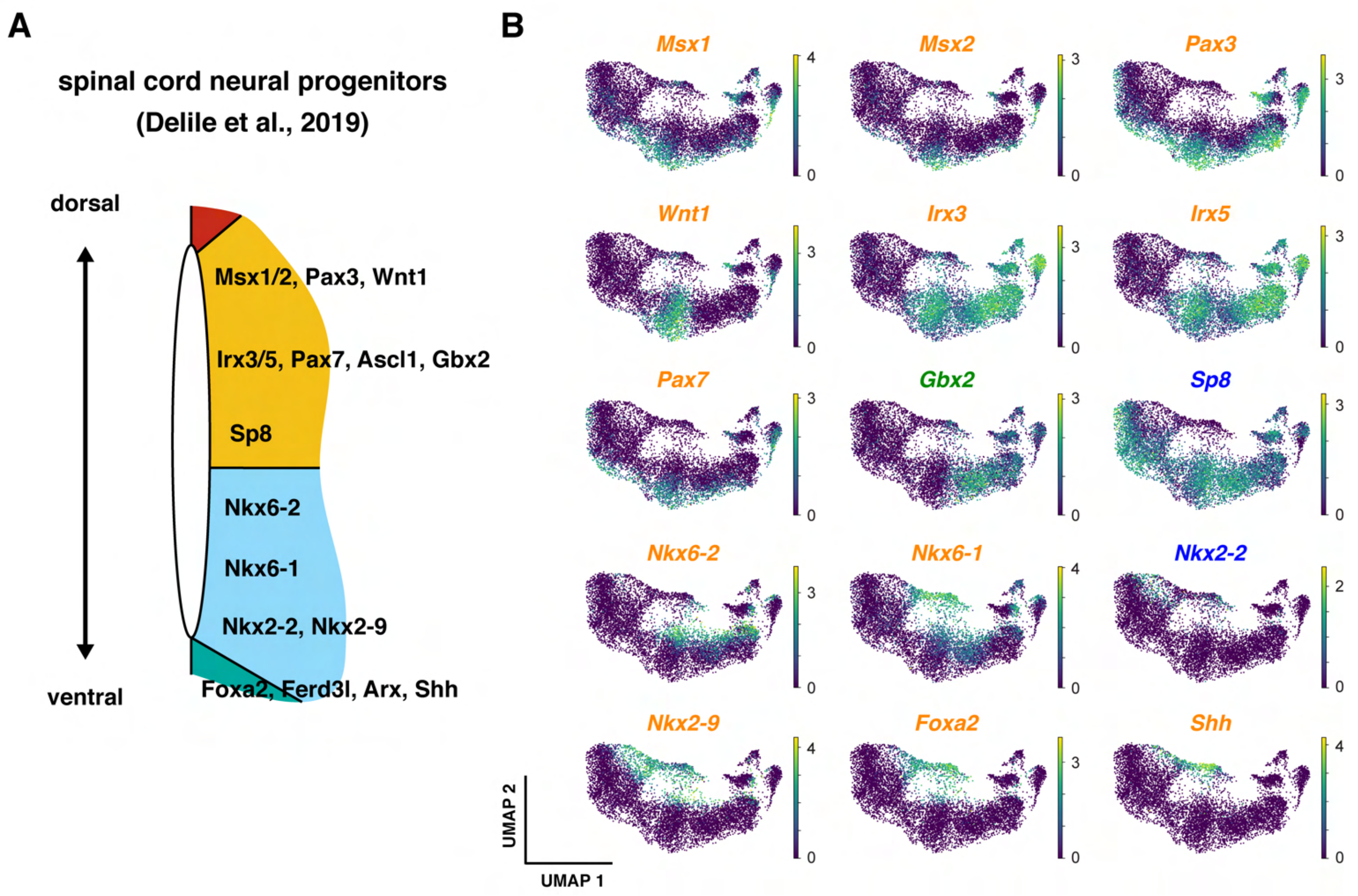

#### Figure 6 - Figure Supplement 1

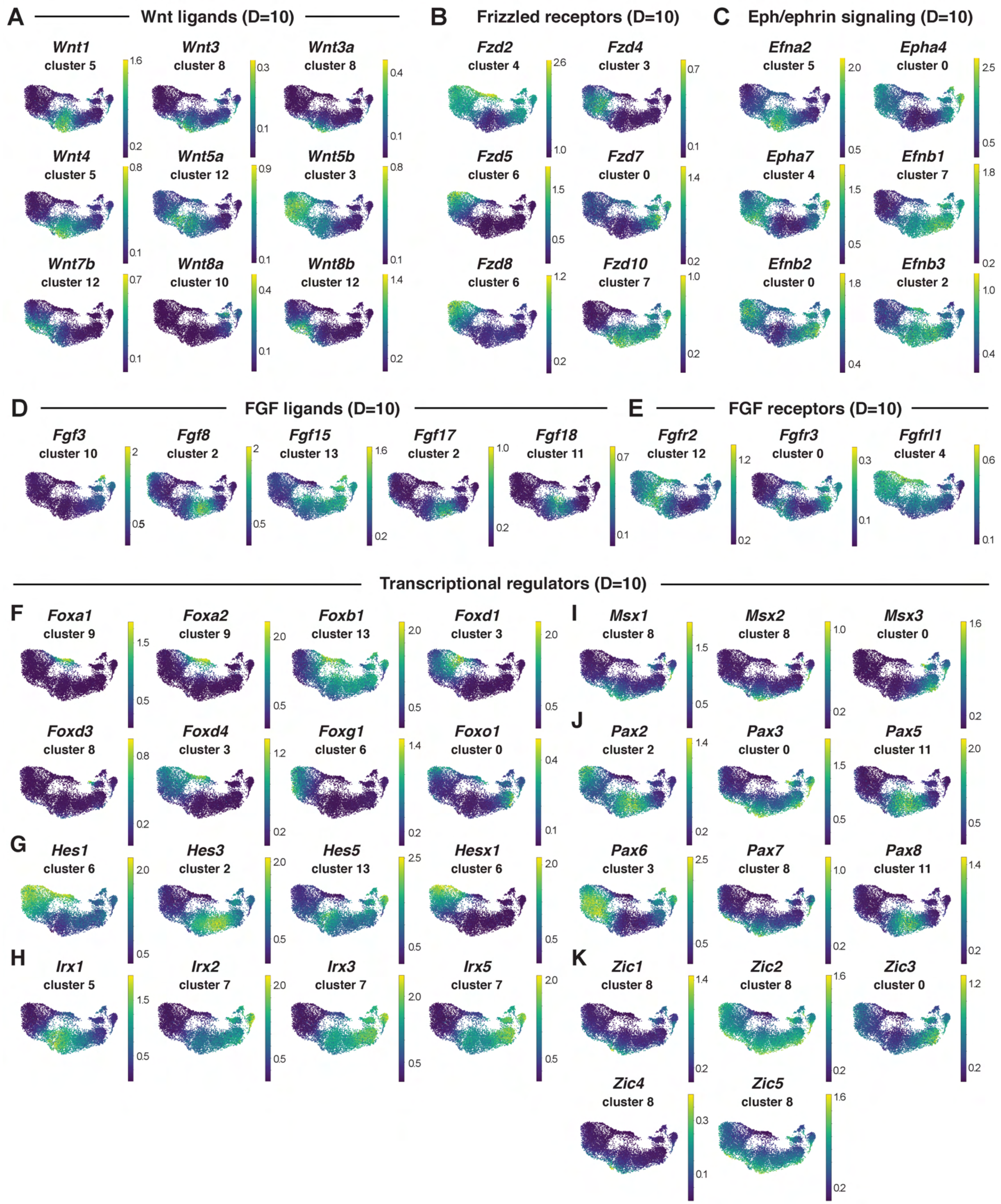
